# Neuroplasticity of language in left-hemisphere stroke: evidence linking subsecond electrophysiology and structural connections

**DOI:** 10.1101/075333

**Authors:** Vitória Piai, Lars Meyer, Nina F. Dronkers, Robert T. Knight

## Abstract

Our understanding of neuroplasticity following stroke is predominantly based on neuroimaging measures that cannot address the subsecond neurodynamics of impaired language processing. We combined behavioral and electrophysiological measures and structural-connectivity estimates to characterize neuroplasticity underlying successful compensation of language abilities after left-hemispheric stroke. We recorded the electroencephalogram from patients with stroke lesions to the left temporal lobe and matched controls during context-driven word retrieval. Participants heard lead-in sentences that either constrained the final word (“He locked the door with the”) or not (“She walked in here with the”). The last word was shown as a picture to be named. We conducted individual-participant analyses and focused on oscillatory power as a subsecond indicator of a brain region's functional neurophysiological computations. All participants named pictures faster following constrained than unconstrained sentences, except for two patients, who had extensive damage to the left temporal lobe. Left-lateralized alpha-beta oscillatory power decreased in controls pre-picture presentation for constrained relative to unconstrained contexts. In patients, the alpha-beta power decreases were observed with the same time course as in controls but were lateralized to the intact right hemisphere. The right lateralization depended on the probability of white-matter connections between the bilateral temporal lobes. The two patients who performed poorly behaviorally showed no alpha-beta power decreases. Our findings suggest that incorporating direct measures of neural activity into investigations of neuroplasticity can provide important neural markers to help predict language recovery, assess the progress of neurorehabilitation, and delineate targets for therapeutic neuromodulation.

## Introduction

Language function is often impaired following stroke to the left hemisphere, compromising quality of life (Cruice et al. 2010; Hilari et al. 2012). Functional compensation by the right hemisphere has been observed in language recovery using functional neuroimaging methods (e.g., Blasi et al. 2002; Hamilton et al. 2011; Turkeltaub et al. 2011). Our current understanding of functional compensation is predominantly based on haemodynamic brain measures. Yet, language processing occurs in the subsecond time scale and language neuroplasticity must also be addressed by electrophysiological measures that track the time course of language function (see for discussion Reid et al. 2016). The present study combined information from behavioral and electrophysiological measures, and estimations of structural connections to assess language neuroplasticity after left-hemisphere strokes. We show that 1) the spectro-temporal profile of brain activity in the intact right hemisphere of stroke patients mirrors the profile of the intact language-dominant left hemisphere in controls, 2) this right-hemisphere functional compensation depends on the integrity of posterior interhemispheric connections, and 3) this neurophysiological compensation mechanism supports a behavioral compensation of language functions.

Electrophysiology provides a direct measure of neuronal activity and the spectro-temporal information in the brain’s activity can provide insights into neuroplasticity (see Reid et al. 2016 for discussion). In the motor domain, the study of neuronal oscillations has led to progress in delineating the changes in bilateral motor cortex associated with functional recovery after stroke (Gandolfi et al. 2015; Rossiter et al. 2014; Shiner et al. 2015). Oscillations measured over the scalp with the electroencephalogram (EEG) reflect the local and long-distance synchronization of large groups of neurons (Nunez and Srinivasan, 2006), which enable processing and communication and are relevant for behavior and disease (Montez et al. 2009; Swann et al. 2015; Uhlhaas et al. 2008). Thus, the spectro-temporal signature of oscillatory activity of a particular brain region provides a key to the region's functional neurophysiological computations. For example, beta desynchronization in the motor cortex is a characteristic fingerprint of the engagement of the motor system in the preparation and execution of both manual (Alegre et al. 2004; Cheyne, 2013) and vocal (Salmelin and Sams, 2002) movements.

The altered spectro-temporal profile of neuronal activity in a particular brain area can indicate whether brain damage has changed the neurophysiological computations in that location (e.g.,Gandolfi et al. 2015; Rossiter et al. 2014; Swann et al. 2015). In the recovery of motor function following stroke, for example, the duration of movement-induced beta-desynchronization and post-movement synchronization were correlated with the patients’ motor function scores (Shiner et al. 2015), indicating that the temporal dimension provided by EEG contributes to the understanding of motor recovery. Similarly, neuroplasticity indexed by spectral and evoked potential changes in working memory and attention has been observed after strokes in lateral frontal cortex (Voytek et al. 2010). The potential of such neuronal oscillations as biomarkers for monitoring neurorehabilitation and recovery of language function is largely unexplored (e.g., Kielar et al. 2016; Meltzer et al. 2013; Nicolo et al., 2015; Spironelli & Angrilli, 2009; Spironelli et al. 2013).

In spontaneous language production, sentence constraint is a major determinant of fluency (Levelt, 1989). By constraint we mean that the evolving conceptual content of the message being conveyed guides access to lexical candidates through spreading activation in the lexico-semantic network, facilitating word retrieval (Griffin and Bock, 1998; Levelt, 1989). Here, we employed a task that comes as close as possible to this naturalistic setting. Participants named pictures following lead-in sentences that either provided a semantic context for the picture (e.g., “He locked the door with the [picture: key]”) or not (“She walked in here with the [picture: key]”). An example of the trial structure is shown in Fig. 1. This task has a number of advantages that makes it ideal for the present investigation. First, the evolving sentence context elicits robust, replicable oscillatory modulations with normal participants localized to the left hemisphere (Piai et al. 2014a, 2015). Given that the context effect is a comparison of two conditions requiring language comprehension and production, it does not suffer from potential problems of either comparing task performance to a rest baseline or comparing two different tasks (e.g., a phonological and a semantic task). Furthermore, the oscillatory activity measured with this task is well understood in relation to memory function: desynchronization of alpha-beta oscillations, reflecting cortical excitation, are a spectral fingerprint of memory retrieval (Hanslmayr et al. 2012).

**Fig. 1.**
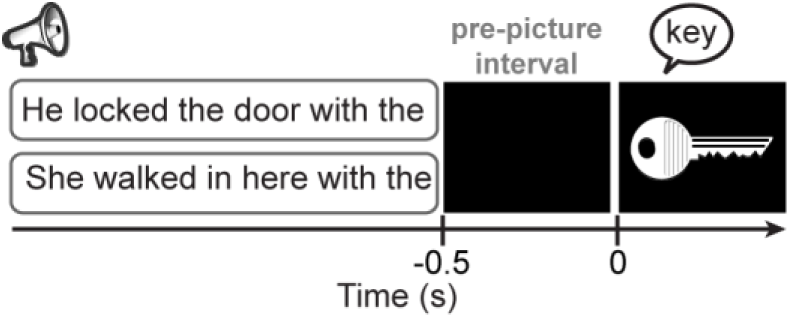
Trial structure. An example of a trial with constrained (upper) and unconstrained (lower) sentence contexts with auditory sentences and visual pictures. Only one sentence was presented at each trial. Sentences started between 3.1 and 3.6 s before picture presentation.

Using this paradigm in a healthy population of younger adults (Piai et al. 2014a, 2015), we have shown that picture naming is faster with constrained than unconstrained sentence contexts (Griffin and Bock, 1998). Across different studies (Piai et al. 2014a, 2015; see also Rommers et al. in press, submitted), left-lateralized alpha-beta desynchronization has been consistently found for constrained relative to unconstrained contexts prior to picture presentation. Confirming the left-lateralized topographies of the alpha-beta desynchronization across studies, the effect has been localized to left angular and supramarginal gyri, left anterior and posterior temporal cortex and left inferior frontal cortex using magnetoencephalography (Piai et al. 2015). These results provided clear evidence that memory retrieval during word production is reflected in alpha-beta desynchronization. Thus, by examining alpha-beta desynchronization prior to picture presentation in this paradigm, we are assessing a well-established marker of context-driven language processes involved in retrieving the name of the picture, which in a healthy population is associated with left temporal and inferior parietal cortices.

White-matter pathways play an important role in recovery of function, both in the language-dominant hemisphere (e.g., Geva et al. 2015), as well as in the contralesional hemisphere (e.g., Forkel et al. 2014). In older adults, the ability to perform a demanding lateralized task through the recruitment of bilateral brain areas seems to depend on the integrity of white-matter connections in the corpus callosum (Davis et al. 2012). However, little is known about the role of interhemispheric connections in the neuroplasticity of language following left-hemisphere strokes.

Six chronic stroke patients with lesions in the left hemisphere (see Table 1 and Fig. S1 for lesion details) and six matched controls participated in the study. Patients’ lesions maximally overlapped in the left middle temporal gyrus, as shown in Fig. 2A. The patients’ language profiles varied but all patients had auditory verbal comprehension and naming abilities that allowed them to perform the task (see Table 2 and Supplementary Table S1). Following previous findings (Piai et al. 2014a, 2015; Rommers et al. in press, submitted), we predicted that controls would show alpha-beta desynchronization prior to picture presentation for constrained relative to unconstrained contexts. For the patients, given our previous demonstration of the involvement of the left temporal cortex in the context effect (Piai et al. 2015), two predictions emerged. Firstly, if the left temporal cortex is a critical structure for benefiting from sentence context in word retrieval, our patients should show little to no facilitation from constrained sentences in picture naming and, accordingly, no alpha-beta power modulations prior to picture onset. Secondly, plasticity in the language network could enable the intact right hemisphere to compensate for the damage in the left hemisphere. If so, we should observe a behavioral facilitation in the picture-naming RTs accompanied by the same spectro-temporal fingerprint of context-driven word retrieval (i.e., alpha-beta desynchronization) but with a spatial configuration switched to the right hemisphere. Finally, we hypothesized that the structural integrity of posterior callosal fibers, directly connecting left and right temporal lobes (Hofer and Frahm, 2006), would facilitate neuroplasticity by engaging right hemispheric structures. Put differently, substantial damage to the posterior corpus callosum would impair the ability of the right hemisphere to compensate for the loss of the language functions subserved by the left hemisphere. For that, we derived the patients’ white-matter disconnection maps (Rojkova et al. 2016) and then calculated the probability of damage to the posterior corpus callosum (i.e., splenium).

**Table 1.** Individual and group lesion volume and percent damage to the left middle temporal gyrus (MTG) and superior temporal gyrus (STG), angular gyrus (AG) and supramarginal gyrus (SMG), and left inferior frontal gyrus (LIFG).

| Patient | Lesion volume<br>(cm<sup>3</sup>) | % STG and MTG | % AG and SMG | % LIFG |
| --- | --- | --- | --- | --- |
| P1 | 18.32 | 26.9 | 7.4 | 0 |
| P2 | 4.51 | 5.6 | 0 | 0 |
| P3 | 93.75 | 62.3 | 48 | 0 |
| P4 | 36.95 | 45.5 | 0 | 0 |
| P5 | 103.17 | 22.7 | 21.7 | 21.4 |
| P6 | 85.82 | 84.5 | 43.1 | 0 |
| Mean (sd) | 57.08 (38.68) | 41.25 (28.82) | 20 (21.4) | 3.57 (8.74) |

**Fig. 2.**
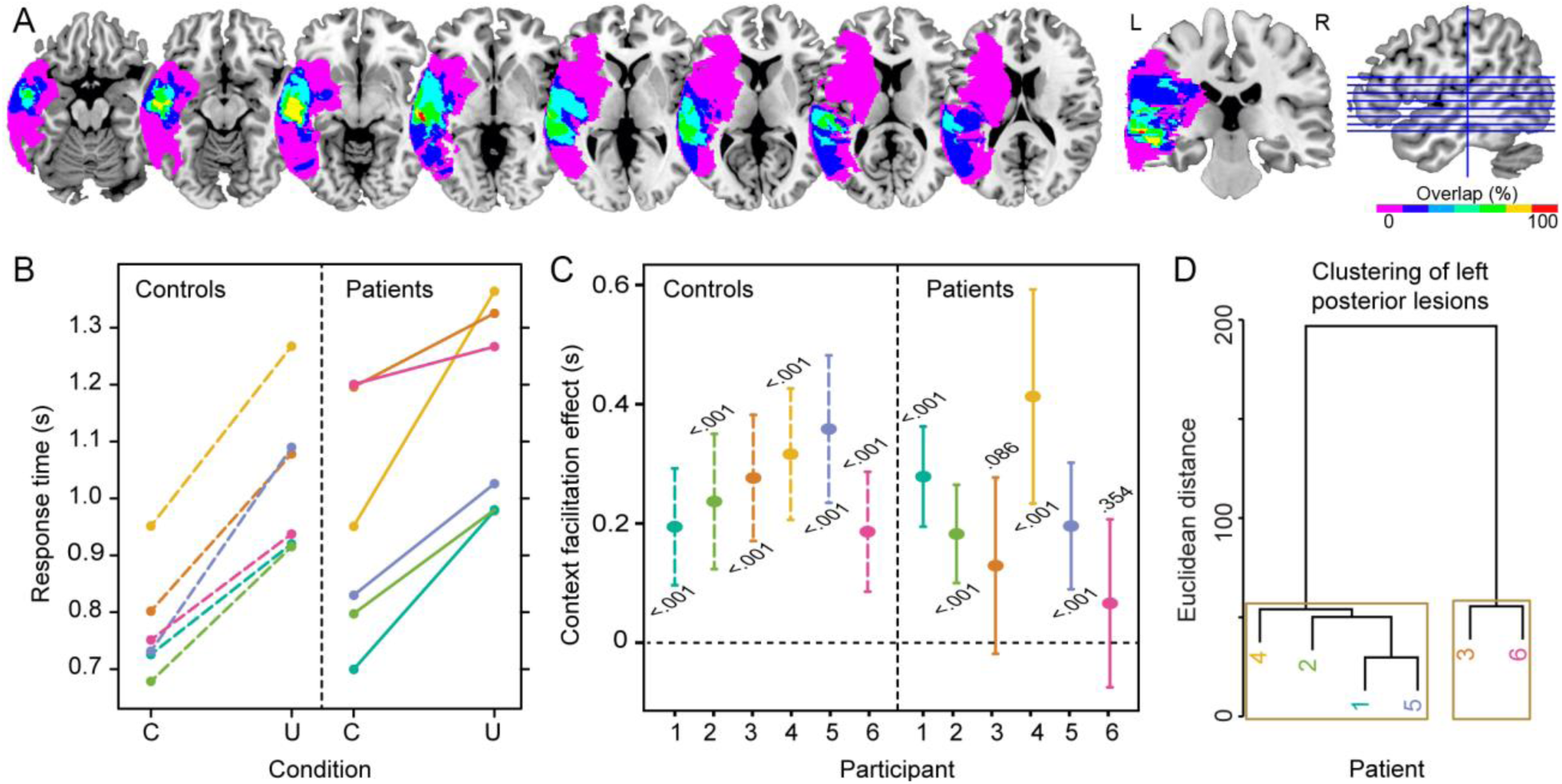
Lesion overlap map and behavioral results. A. Lesion overlap map of the patients. The color scale indicates the amount of overlap in lesion locations, with magenta indicating that only one patient had a lesion in that particular region (i.e., 0% overlap). The maximum overlap (100%), shown in red (4^th^ axial slice from left to right and the coronal slice), indicates that all patients had a lesion in the middle temporal gyrus. B. Mean picture naming times for each participant and condition (C = constrained; U = unconstrained). C. Context facilitation effect with 95% confidence intervals and significance (p value) for each participant. The color correspondence between controls and patients indicates the link between each patient and the matched control participant. D. Dendogram of the left posterior lesion clusters over superior temporal, middle temporal, angular, and supramarginal gyri. Brown boxes indicate the two significant clusters.

**Table 2.** Language testing data from the Western Aphasia Battery (WAB) and time elapsed between stroke date (MPO = months post onset) and WAB testing, and between stroke date and the present EEG experiment. Naming = WAB Naming and Word Finding score (maximum = 10). Comprehension = WAB Auditory Verbal Comprehension score (maximum = 10). Sequential commands = WAB comprehension subtest (maximum = 80). Aphasia Quotient (AQ, maximum = 100). WNL = within normal limit.

| Patient | Aphasia type | AQ | Naming | Comprehension | Sequential commands | MPO at WAB | MPO at EEG |
| --- | --- | --- | --- | --- | --- | --- | --- |
| P1 | NA* | NA* | NA* | NA* | NA* | NA* | 114 |
| P2 | WNL | 99.6 | 10 | 10 | 80 | 104 | 121 |
| P3 | Conduction | 77.9 | 8.6 | 8.55 | 58 | 16 | 23 |
| P4 | WNL | 94 | 8.6 | 10 | 80 | 222 | 230 |
| P5 | Anomic | 87.8 | 8.3 | 8.15 | 43 | 47 | 72 |
| P6 | Anomic | 92.9 | 9.5 | 9.55 | 72 | 290 | 310 |

\*P1 was not assessed on the WAB as he was not aphasic, see Methods section for language profile.

### Experimental Procedures

The study protocol was approved by the University of California, Berkeley Committee for Protection of Human Subjects, following the declaration of Helsinki. All participants gave written informed consent after the nature of the study was explained and received monetary compensation for their participation.

### Participants

Six patients with lesions to the left lateral temporal cortex participated (one female; median age = 68, mean = 65, sd = 10, range = 50-74; mean years of education = 17). All patients were tested at least 23 months post stroke and were pre-morbidly right handed. Information on the patients' lesions and language ability are shown in Tables 1 and 2.

Patients’ language abilities were assessed with the Western Aphasia Battery (WAB, Kertesz, 1982). The language profiles for five of these patients are shown in Table 2, together with the time elapsed between patients’ stroke date and WAB assessment, and stroke date and EEG testing. Three of the six patients had good language abilities (P1, P2, and P4), with performance within normal limits. We note that, although P1 was not assessed on the WAB, he continued performing his occupation without problems, which included academic teaching amongst other tasks. Thus, we are confident that this patient would have been classified as within normal limits by the WAB. Two patients were classified as anomic (P5 and P6), characterized by normal auditory verbal comprehension and repetition, but a relatively impaired word finding ability when speaking. Finally, one patient was classified as having conduction aphasia (P3), characterized by normal auditory verbal comprehension, but relatively impaired repetition and word-finding abilities. Additionally, six controls participated, each matched closely to each patient for gender (one female), age (matched within ±4 years; median age = 68, mean = 66, sd = 9, range = 54-74, Mann–Whitney U = 16, *p* = .809), and years of education (matched within ±2 years; mean years of education = 18, Mann–Whitney U = 12, *p* = .314). None of the patients or control participants had a history of psychiatric disturbances, substance abuse, medical complications, multiple neurological events, or dementia. All participants were native speakers of American English.

## Materials

Fifty-one words were chosen for which we selected colored public-domain clipart images from the internet. For each word, two sentences were created for which the target word was the last word of the sentence, presented as a picture. All 102 sentences had six syllables. The sentences formed two different conditions. In the unconstrained condition, no specific word was expected as the final word of the sentence whereas for the constrained condition, the target word was highly expected. For each target word, the associated sentences had the same two last words. A pre-test was used to verify that the sentences differed in the degree of expectancy for the final word as a function of context (i.e., cloze probability; Taylor, 1953). Fourteen participants were presented with the sentences up to the penultimate word of the sentence, excluding the final word. Cloze probability was calculated as the proportion of participants who used the target picture name as their completion (constrained mean cloze probability = 0.83; unconstrained mean cloze probability = 0.04, *p* < 0.001). The sentences in the unconstrained condition did not have a high cloze probability for any word.

The sentences were spoken by a female native speaker of American English and recorded in a soundproof booth (sampling frequency = 44.1 kHz) and subsequently normalized to 77 dB sound-pressure level. The sentences were spoken at a regular pace, guided by a metronome set at 127 bpm, yielding 2.11 syllables per second on average. The mean duration of the sentences was equal across conditions (context: mean = 2.8 sec, sd = 0.10, range = 2.6 – 3.1; no-context: mean = 2.8 sec, sd = 0.13, range = 2.6 – 3.1, paired-samples *t*(50) < 1). Pitch contours did not differ between conditions (tested as mean pitch over three segments of the speech sound), all *p*s > 0.05.

### Procedure

Stimulus presentation and response recording was controlled by Presentation (Neurobehavioral Systems, Albany, CA). After EEG preparation, participants were brought individually to an electrically-shielded, sound-attenuated, dimly-lit booth. A practice session was performed first, in which the participants trained the naming of pictures without collateral blinking. A fixation cross was presented for 1.5 s, followed by the picture, which remained on the screen for 1.5 s. Then, a black screen appeared for 1 s followed by three asterisks for 1.5 s, indicating that participants could blink. The same asterisk screen was used during the task proper to help participants postpone their blinking until after naming the pictures. The sentences were presented via stereo loudspeakers, after participants confirmed that the volume was optimal for comprehension (no adjustments had to be made, so volume remained equal for all 12 participants). A trial began with a fixation cross, displayed continuously during auditory sentence playback. After 1 s, the sentence was presented. After sentence offset, the fixation cross remained on the screen for another 0.5 s before the picture was displayed for 2 s. Then, the three asterisks appeared to indicate blinking for a variable interval between 1.2 and 1.9 s. An example of an experimental trial is given in Fig. 1.

### EEG acquisition

EEG was recorded from 64 Ag/AgCl pre-amplified scalp electrodes (BIOSEMI, Amsterdam, Netherlands) mounted in an elastic cap according to the extended 10-20 system. EEG was sampled at 1024 Hz. The electrooculogram was recorded horizontally from electrodes placed on the left and right temples and vertically from Fp1 and the electrode positioned below the left eye. Surface electromyogram was recorded from the orbicularis oris muscle with two electrodes placed on the left upper and right lower corner of the mouth.

### MRI acquisition

Anatomical magnetic-resonance images of the patients were acquired in a separate session using a Siemens Magnetom Verio 3T MRI scanner (Siemens, Germany), equipped with a 12-channel head coil. A T1-weighted 3D MPRAGE imaging sequence (Brant-Zawadzki et al.1992) was applied (parameters: TR /TE= 3000/3.54 ms, FOV = 240 mm, flip angle= 8o, resolution = 256 × 256 × 212, voxels = 0.94 × 0.94 × 1.3 mm, 5 min acquisition time).

### Behavioral analysis

The experimenter monitored participants’ naming responses online for errors (i.e., disfluencies, omissions, or unrelated responses). Near-synonyms were considered correct (e.g., “mug” and “cup”). Errors were few (1.3% for controls and 5.2% for patients) and excluded from further analyses. An overview of the types of errors made by the patients is shown in Supplementary Table S1. Response time (RT) analyses comprised correct trials only. Response times were calculated manually using the speech waveform editor Praat (Boersma and Weenink, 2013) before the trials were separated by condition. Statistical analyses of the RTs were conducted using R (R Development Core Team, 2014).

Participants’ mean RTs were computed for each condition. Non-parametric tests were used to evaluate the behavioral effects at an alpha level of 0.0167 (two-tailed). The overall context effect (constrained vs unconstrained) was assessed with a Wilcoxon signed-rank test. The overall group effect (controls versus patients) and the difference in context effect between the two groups (i.e., the interaction) were assessed with Mann-Whitney-Wilcoxon tests. Moreover, the context effect was also evaluated within each participant using independent-samples *t*-test across trials. In the Supplement (Fig. S2), we report the results when the context effect was evaluated within each participant using dependent-samples t-tests across trials.

### Lesion analysis

Lesions were drawn on patients’ structural magnetic resonance imaging scans by a trained technician and confirmed by a neurologist (RTK). For EEG source localization, lesions were left in their native space. For the other analyses, lesions were normalized to the MNI template. A hierarchical clustering analysis was run on the percent damage of left middle temporal gyrus (MTG), superior temporal gyrus (STG), angular gyrus (AG), and supramarginal gyrus (SMG) using the Euclidean distance and the Ward criterion. Statistical significance was assessed using multiscale bootstrap resampling with 1,000 bootstraps (Suzuki and Shimodaira, 2006). We also mapped the lesion from each patient onto tractography reconstructions of white-matter pathways obtained from a group of healthy controls (Rojkova et al. 2016). The severity of the posterior interhemispheric disconnection was calculated individually from the probability of the splenium tract to be disconnected, thresholded at > 80% probability, using Tractotron software as part of the BCBtoolkit (Thiebaut de Schotten et al. 2014, http://www.brainconnectivitybehaviour.eu). This stringent threshold was chosen to increase the reliability of our results. In addition, for each patient, we derived a normalized measure of lesioned tissue (in percentage) for grey matter (GM) and white matter (WM) separately (cf. Meyer et al. 2014). For that, we masked a canonical GM and WM tissue probability map (SPM12 software package, Wellcome Trust Centre for Neuroimaging Department, University College, London, UK) with the patients’ MNI-normalized lesion volumes. Then, we counted the damaged voxels that fell into GM and WM, respectively (thresholded at > 80% probability in analogy to the splenium analysis; see above). Finally, to normalize for the differences in size between GM and WM structures, the total counts of damaged voxels in GM and WM were divided by the total number of GM and WM voxels in the respective whole-brain tissue probability maps (thresholded at 80% probability).

### EEG analysis

The analyses were performed using FieldTrip version 20150927 (Oostenveld et al. 2011) in MatlabR2014a. All trials excluded from the RT analysis were also excluded from EEG analysis. Each electrode was re-referenced off-line to averaged mastoids. The data were high-pass filtered at 0.16 Hz (FieldTrip default filter settings) and segmented into epochs time-locked to picture presentation, from 800 ms pre-picture onset to 300 ms post-picture onset. All epochs were inspected individually for artifacts such as eye-movements, blinks, muscle activity, and electrode drifting. Eight peripheral channels (T7, T8, TP7, TP8, F7, F8, FT7, FT8) were excessively noisy (variance > 4 millivolt) in the majority of participants, and were therefore removed from analyses. All but two participants (P6 and C4) moved their eyes (saccades and eye blinks) during the blinking interval in most of the trials. For these participants, trials with eye movements were excluded. For P6 and C4, independent component analysis was used to correct for eye movements (Jung et al. 2000, as implemented in FieldTrip). One clear eye-movement component was removed for each of the two participants (Figure S3). On average, error-and artifact-free trials comprised 44 trials per condition for patients (no difference in trial numbers between conditions, Wilcoxon signed rank *p* = 0.854) and 47 trials for controls (no difference in trial numbers between conditions, Wilcoxon signed rank *p* = 0.202). No difference was found between the number of available trials for patients and controls, Mann-Whitney-Wilcoxon *p* = 0.108). Time-frequency representations were calculated from the cleaned EEG segments. Power was calculated with a modified spectrogram approach, at frequencies ranging from 8 to 30 Hz (following findings from Piai et al. 2014a, 2015). We used an adaptive sliding time window of three cycles’ length (e.g., the window was 300 ms long at 10 Hz, see for a similar approach Haegens et al 2011; Piai et al. 2014a, 2014b). This window was advanced in steps of 10 ms in the time dimension and in steps of 1 Hz in the frequency dimension. The data in each window was multiplied with a Hanning taper, and the Fourier transform was taken from the tapered signal. For each group (patients and controls), we compared the time-frequency representations in the two conditions using a non-parametric cluster-based permutation test (Maris and Oostenveld, 2007) based on dependent-samples t-tests. For the interaction contrast of group and condition, the difference in power between the two conditions in each group was compared to each other using independent-samples *t*-tests. The statistical analyses thus comprised the alpha and beta bands (8 to 30 Hz), all channels, and all time points, and were assessed at an alpha level of 0.05 (two-tailed).

### Source-level analysis

For source-level analysis, we focused on the time-frequency window of the significant sensor-level effect (see Results). From each patient's T1-weighted scan, we generated an individual finite-element headmodel (Wolters et al. 2004). Using the SPM12 software package (Wellcome Trust Centre for Neuroimaging Department, University College, London, UK) and FSL (FMRIB, University of Oxford, UK), each patient's image was segmented into gray matter, white matter, cerebro-spinal fluid, skull, and scalp. The manually delineated lesion segment was combined with the cerebro-spinal fluid segment, and hexahedral meshes were generated for all tissue types. Standard tissue conductivities were used (Vorwerk et al. 2014). Template electrode positions were aligned to the FEM's fiducial axes and then projected to the scalp mesh. A template 10-mm-spaced 3-D source grid was generated inside the gray matter of the MNI template and warped to each individual FEM for beamforming. An individual Dynamic Imaging of Coherent Sources (Gross et al. 2001) beamformer was employed on the time-frequency data across conditions, generating for each grid point a common spatial filter. The common filter was then applied to the single-trial data from the individual conditions. Patients’ individual source localization of the alpha-beta effect of interest was performed with non-parametric cluster-based permutation tests based on independent-samples *t*-tests between conditions, over trials. Using the Automated Anatomical Labeling atlas (Tzourio-Mazoyer et al. 2002), we extracted and averaged the *t* values from the left and right lateral temporal cortex and inferior parietal lobe (i.e., bilateral AG and SMG). Using *t* values instead of averaged power gives us a more robust within-participant measure. Given that standard statistics on single-trial source-level results do not have enough power to yield significance on the individual-level, we used a threshold of *t =*-1 to indicate the reliability of the context effect in each patient. The relationship between reliable power decreases (i.e., *t* values exceeding −1) in the temporal and inferior parietal lobes and patients' behavioral effects was examined with a Fisher's exact test in the form of a contingency table, with a mid-p correction (Lancaster, 1961). The relationship between damaged tissue and power decreases was assessed with non-parametric Spearman's rank correlation coefficients after averaging all source-level *t*-values in each hemisphere separately.

## Results

### Sentential context facilitates word retrieval in both patients and controls

As Fig. 2B shows, all participants were faster in the constrained than in the unconstrained condition. Patients' mean RTs were 946 ms (sd = 211) for the constrained and 1157 ms (sd = 181) for the unconstrained condition. Controls’ mean RTs were 773 ms (sd = 96) for the constrained and 1035 ms (sd = 139) ms for the unconstrained condition. Responses in the constrained condition were faster overall than in the unconstrained condition (Wilcoxon signed rank = 1, *z* = −3.30, *p* < .001, 95% CI [−272 −129]). Patients’ and controls overall RTs did not differ significantly from each other (Mann–Whitney U = 45, z = −1.52, *p* = .128, 95% CI [-313 36]). There was no statistically significant difference in the magnitude of the context effect between groups (Mann–Whitney U = 19, z = −0.08, *p* = .937, 95% CI [-121 203]).

Fig. 2C shows the context facilitation effect for each participant individually (and their respective 95% confidence intervals and within-participant significance value, see also Fig. S2). Within-participant statistical analyses over trials indicated that four of six patients (P1, P2, P4, P5) were faster on constrained trials, indicating a robust within-participant behavioral facilitation effect.

### Large lesions to left temporal-parietal cortex affect contextual processing

Using MEG, we have previously shown that the contextual alpha-beta desynchronization effect localizes to the left MTG, STG, AG, and SMG (Piai et al. 2015). Based on these findings we entered the percentage of left MTG, STG, AG, and SMG damage separately in a hierarchical clustering analysis in order to group patients according to their lesion profile. The results indicated two distinct clusters (Patients 3 and 6 vs. Patients 1, 2, 4, 5; multiscale bootstrap resampling *p*s < 0.01), as shown in Fig. 2D. Patients 3 and 6 have the most volume loss in left MTG, STG, AG, and SMG (see Table 1 and Supplementary Fig. S1 for individual lesions), and show a reduced behavioral facilitation with no significant context effect over trials (see Fig. 2C).

### Sentential context modulates pre-picture alpha-beta power

For both patients and controls, alpha-beta power decreased between 20% and 25% prior to picture presentation. We used cluster-based permutation tests (Maris and Oostenveld, 2007), which effectively control the false alarm rate, in an uninformed manner (i.e., without a-priori constraints on time points, frequency points, and channels assessed). Statistically significant differences were found between the context conditions for both controls (*p <* 0.001) and patients (*p <* 0.001) that could be attributed to a spatio-spectro-temporal cluster of power decreases in each group. These clusters were detected between 8 and 25 Hz and −0.4 to 0 s in both groups, but they were spatially distinct, as shown in Fig. 3. Whereas the controls showed a left-lateralized alpha-beta desynchronization, replicating previous studies (Piai et al. 2014a, 2015), the left temporal patients showed a right-lateralized desynchronization. The interaction between relative power changes and group did not yield any clusters. This finding is to be expected because the degree of power decreases, as well as the spectro-temporal profile of the clusters in the two groups is similar. Thus, rather than a quantitative difference in the amount and in the spectro-temporal signature of power decreases, the difference between patients and controls is spatial in nature. To support this interpretation, we calculated a laterality index for patients and controls individually by subtracting the averaged *t*-value of all left-scalp channels from all right-scalp channels. Controls’ alpha-beta scalp power decreases were left-lateralized in all participants (mean = –0.170, median = –0.162, sd = 0.107) whereas in patients, power decreases were right-lateralized (mean = 0.142, median = 0.094, sd = 0.164). Lateralization differed significantly between controls and patients (Mann–Whitney U = 36, *z* = −3.07, *p* = .002, 95% CI [0.099 0.517]). Figure S4 shows the scalp topography of the alpha-beta desynchronization effect of each control participant, as well as the individual laterality indices.

**Fig. 3.**
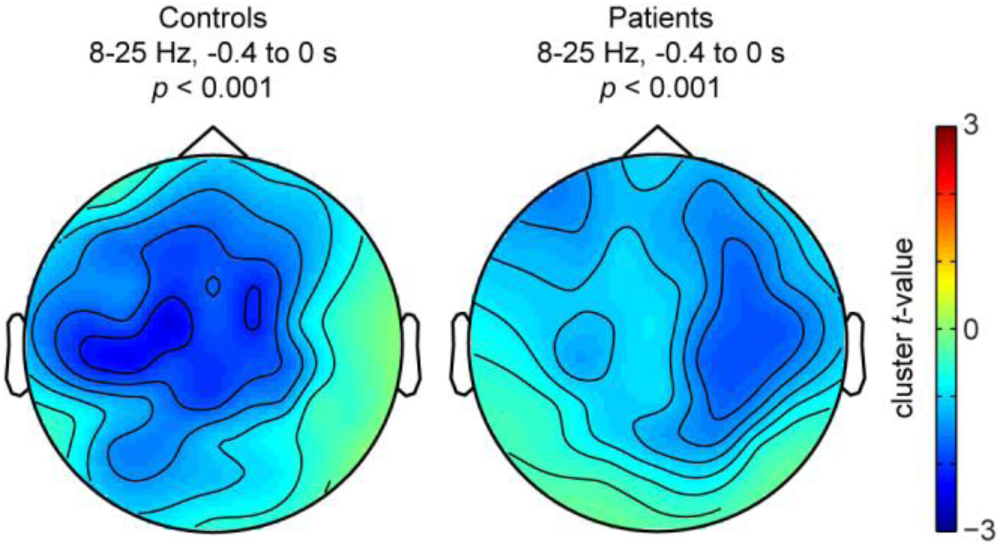
Scalp context effect. Topographical distribution of the significant cluster associated with the relative power decreases between 8 and 25 Hz and −0.4 to 0 s for each group and corresponding p values.

### Right-hemisphere desynchronization in patients

To follow up on these topographical differences in the context effect between patients and controls, we source-localized patients’ individual context effect using frequency-domain beamformers across the significant time-frequency window (see previous section) from −0.4 to 0 s and from 8 to 25 Hz (i.e., the significant cluster). The individual source results of the power decreases are shown in Fig. 4A. The amount of power decreases in the left and right temporal and inferior parietal lobes is summarized in Fig. 4B. The two patients with the least robust power decreases in the temporal and inferior parietal lobes (i.e., *t* values not exceeding −1) were also the two patients with the weakest behavioral effects (patients 3 and 6), *p* = 0.033. Amongst the patients with more robust behavioral and oscillatory context effects (1, 2, 4, and 5), power decreases were statistically stronger in the right than in the left hemisphere for patients 1, 2, and 4. The only patients with *left* temporal and inferior parietal power decreases that exceeded the −1 *t*-value threshold were the two with no posterior MTG involvement (patients 2 and 5, green in Fig. 4C). By contrast, the four patients with *no* left-temporal and inferior parietal power decreases exceeding the −1 *t*-value threshold have lesions that overlap 100% in posterior MTG (pink in Fig. 4C).

**Fig. 4.**
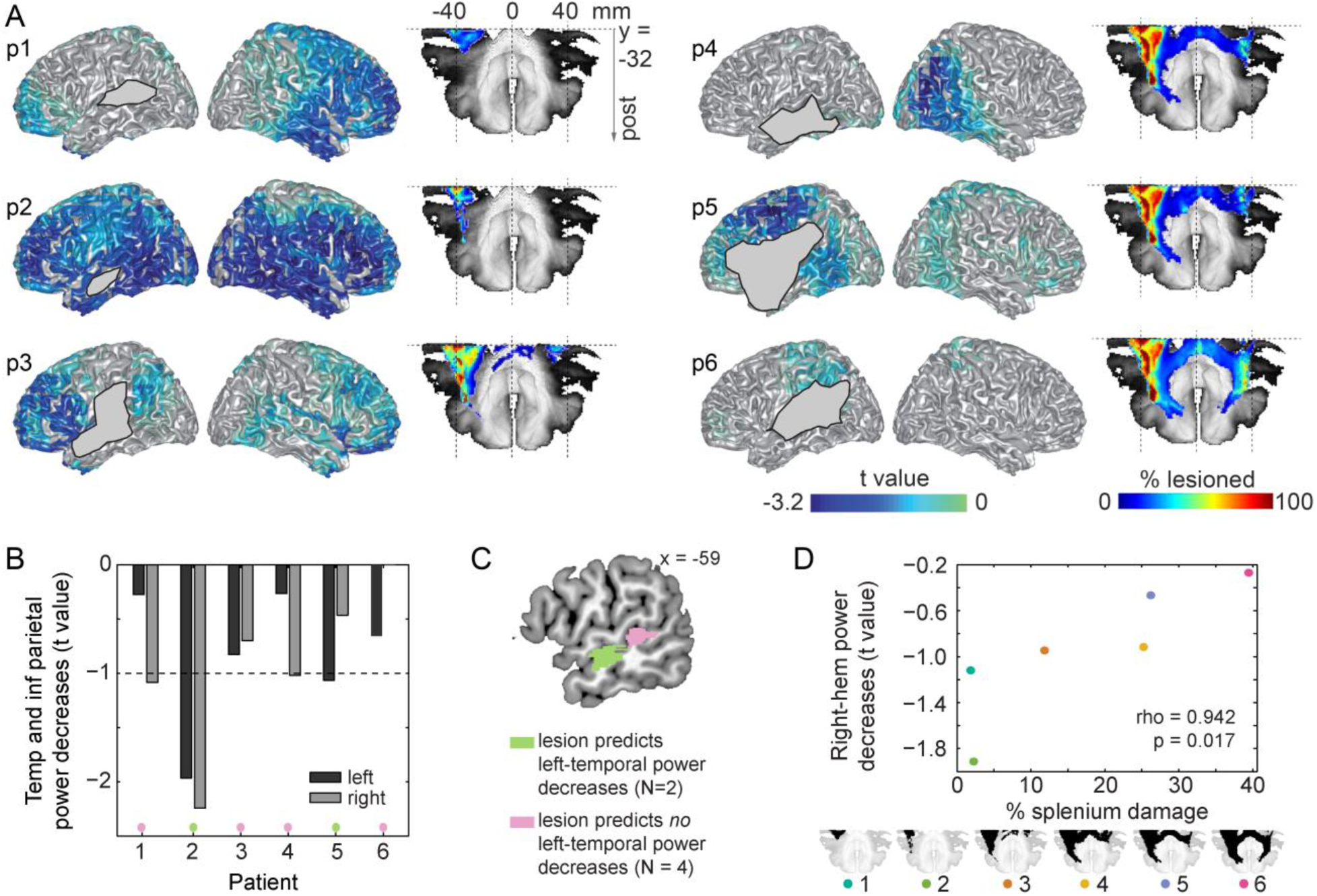
Source context effect and white matter pathways. A. Individual source localization of the power decreases (expressed in t values) between 8 and 25 Hz for all six patients (numbered p1-p6). Lesion masks display where lesioned voxels intersect with the cortical mesh. For patients’ accurate lesion delineation, see Supplementary Fig. S1. Splenium tract and individual disconnected probability maps of the splenium (in *%* lesioned voxels) are shown to the right of each individual source map. B. Mean power decreases in the left (dark grey) and right (light grey) temporal and inferior parietal lobes for each patient. Power decreases are expressed in *t* values given that this is a more robust within-participant measure than mean power is. C. Lesion overlap of the two patients with left temporal and inferior parietal lobe power decreases exceeding the −1 *t*-value threshold (green) and lesion overlap of the four patients with left temporal and inferior parietal lobe power decreases not exceeding the −1 *t*-value threshold (pink). D. Scatterplot of the percentage splenium damage and power decreases in the right hemisphere. For each patient, the splenium is shown in black at the bottom. Post = posterior; temp = temporal; inf = inferior; hem = hemisphere.

Using the normalized GM- and WM-lesion measure, we observed that overall WM damage in our patients was significantly larger than overall GM damage (Wilcoxon signed rank *p* = 0.031). Posterior callosal fibers in the splenium, extending into the tapetum, directly connect the temporal lobes across hemispheres (Hofer and Frahm, 2006; see alsoTurken and Dronkers, 2011). Therefore, we investigated the relationship between the probability of splenium integrity and hemispheric lateralization of the observed alpha-beta power decreases. A sagittal slice of the splenium disconnection probability map of each patient is shown in Fig. 4A. The probability of splenium damage was correlated with alpha-beta power decreases in the right hemisphere (*rho* = 0.942, *p* = 0.017, Fig. 4D), as well as with alpha-beta power decreases in the right temporal and inferior parietal lobes (*rho* = 0.886, *p* = 0.033).

We also assessed the specificity of this relationship by examining the correlation between right-hemisphere power decreases and other structural measures. Figure S5 shows the relationships between the structural variables and the right-hemisphere power decreases. Total lesion volume did not predict either left- or right-hemisphere power decreases (*p*s > 0.136), nor left-or right-temporal lobe and inferior parietal power decreases (*p*s > 0.297). Moreover, the percentage of damage to the superior temporal, middle temporal, angular, and supramarginal gyri did not predict right-hemisphere power decreases (*p*s > 0.175). Finally, right-hemisphere power decreases did not correlate with overall GM damage (*rho* = .543, *p* = .297). Right-hemisphere power decreases had some relationship to overall WM damage, but the correlation was not significant (*rho* = .829, *p* = .058). To further support this finding, we ran a linear regression model with the probability of splenium damage as a predictor and another model with overall WM damage as a predictor of the right-hemisphere power decreases. Overall WM damage did not predict right-hemisphere power decreases (beta = 0.08, *t* = 1.38, df = [1,4], *p* = 0.241, residual standard error = 0.529, adjusted R^2^ = 0.152). Splenium damage did predict right-hemisphere power decreases (beta = 0.03, *t* = 3.24, df = [1,4], *p* = 0.032, residual standard error = 0.338, adjusted R^2^ = 0.655). Thus, the splenium provided the strongest and most robust relationship between structure and functional lateralization, as measured by right-hemisphere alpha-beta power decreases.

## Discussion

In the present study, we used a novel approach to investigate neuroplasticity of language function integrating data from patients’ behavioral performance, electrophysiology, and estimations of structural connectivity. Our results provide three main findings. Firstly, we show a causal link between context-driven word retrieval and alpha-beta desynchronization in the left temporal-inferior parietal lobe. Secondly, we show that following stroke, the right hemisphere can perform similar neuronal computations as the left hemisphere, as evidenced by a comparable spectro-temporal profile in the controls’ left hemisphere and the patients’ right hemisphere. Thirdly, we provide findings suggesting a potential role for posterior transcallosal white-matter connections via the splenium in right-hemisphere language compensation following left hemisphere brain injury.

The reliable right-hemisphere power decreases had a similar spectro-temporal signature as what is observed in the left hemisphere of normal participants (Piai et al. 2014a, 2015, and the present results). This finding suggests that the right hemisphere can perform similar neuronal computations as the left hemisphere performs. Moreover, the three patients with prominent right-hemisphere power decreases showed no behavioral deficits in our task relative to the neurotypical controls. Our findings of right-hemisphere activity and its spectro-temporal signature were obtained in a data-driven manner, with a rigorous statistical test that controls the false-alarm rate in light of the large number of time-frequency points and channels examined (Maris and Ostenveld, 2007). Thus, these findings were not determined by biases in data preselection, but emerged as the strongest feature of our data.

It is debated whether right-hemisphere activity observed in recovery from aphasia is beneficial or maladaptive (e.g., Hamilton et al. 2011; Turkeltaub et al. 2011). Our study was not designed to specifically address this question. However, the patients’ right-hemisphere activity was associated with a context facilitation effect of the same magnitude as the controls’. Moreover, the two patients with slower naming RTs and weak context facilitation effects had no clear alpha-beta modulations in either hemisphere. Together, these results suggest a supportive role for the right hemisphere in context-driven word retrieval. This suggestion is in line with re-emerging evidence on the supportive role of the right hemisphere in the recovery of aphasia (e.g., Barrett & Hamilton, 2016; see also Kielar et al. 2016 for evidence from alpha-beta oscillations).

The two patients with extensive damage to temporal and inferior parietal lobes were impaired in using linguistic context to guide word retrieval in language production. These patients also showed reduced alpha-beta desynchronization in their EEG. Corroborating this finding, a lesion in the posterior MTG abolished the alpha-beta desynchronization in the left temporal-inferior parietal lobe. These findings indicate a causal link between context-driven word retrieval and alpha-beta desynchronization in the left temporal-inferior parietal lobe.

Our findings provide novel insights into a potential mechanism linking interhemispheric structural connectivity and right-hemisphere functional compensation. The probability of splenium damage was a strong predictor of right-hemisphere alpha-beta desynchronization, whereas damage to other structures, including lesion volume, did not predict any oscillatory effects. The overall WM-damage measure had the second strongest correlation with the right-hemisphere power decreases, although this correlation was not significant. This finding was confirmed by the regression analyses, which indicated that splenium damage was a significant predictor of right-hemisphere power decreases, explaining 66% of the variance, whereas overall WM-damage was not a significant predictor, explaining only 15% of the variance. Thus, we conclude that splenium damage was the best predictor of right-hemisphere functional lateralization in our study.

Some limitations of the present study are worth mentioning. Firstly, our sample size included six patients, heterogeneous with respect to their language deficits. We note however that the study was conducted on a well-delineated target population with lesions consistently affecting the temporal lobe and, in particular, the middle temporal gryus. Although previous studies examining oscillations included more patients, the lesion overlap in the temporal lobe in those studies may not have exceeded the overlap in our sample of six patients (e.g., Kielar et al. 2016; Meltzer et al. 2013; Spironelli & Angrilli, 2009; Spironelli et al. 2013). Moreover, we took advantage of the heterogeneity and performed individual-participant statistical analyses to be able to explain the patterns of brain-behavior relationships observed. Despite the small sample size, all behavioral and electrophysiological effects were sufficiently powered with a sample size of N = 6 at an alpha level of .05 with .90 power (see Supplement). Secondly, most errors in EEG source localization emerge when template head models are used rather than head models based on participants’ native MRIs (Acar and Makeig, 2013). Importantly, this is not an issue for our source results given that we used the patients’ native MRIs and explicitly modelled the lesioned volumes. We could not perform source localization of the context effect in the controls because their MRIs were not available. However, the scalp effect in the controls replicates the left-dominant topography of previous studies (Piai et al. 2014a, 2015) and our scalp laterality index indicated that the effect was left lateralized in all six control participants. Thirdly, we could not measure interhemispheric structural connectivity directly but instead estimated it through the patients’ MRIs. Despite this limitation, our results suggest a strong relationship between probability of interhemispheric structural connections and right-hemisphere functional compensation. This observation opens new avenues for future research.

The question can be asked whether the relatively weak context effect in patients 3 and 6 reflects poor context-driven word retrieval, emerging at the level of spreading activation through the lexical-semantic network, or rather a deficit in sentence comprehension. We argue against this latter account for a number of reasons. Firstly, all patients had high accuracy in picture naming in our task (see Supplementary Table S1). Secondly, patient 6, who had the weakest context effect, scores high on comprehension on the WAB aphasia battery (95% accuracy, see Table 2). Patient 6 also scores high (90% accuracy) on a particular comprehension subtask (i.e., sequential commands), which requires patients to comprehend sentences that are more complex than the six-syllables sentences used in our experiment (e.g., “Point with the pen to the book”). Patient 5, by contrast, scored relatively low in this subtask (i.e., 54%), but showed a strong and reliable context facilitation effect. Thus, a deficit in sentence comprehension alone cannot explain the context effect in our patients.

EEG, and oscillatory activity in particular, provide a biologically meaningful, direct measure of neuronal activity that can help us understand neuroplasticity. The role of alpha-beta desynchronization is well studied. For word comprehension and production, previous studies have indicated that alpha-beta desynchronization is an index of word retrieval (e.g., Brennan et al. 2014; Mellem et al. 2012; Piai et al. 2015). In the aphasia literature, however, this neural signature is largely unexplored (Kielar et al. 2016; Meltzer et al. 2013; Spironelli et al. 2013). Spironelli et al. (2013) had participants with non-fluent aphasia perform semantic, phonological, and orthographic tasks in which a button-press response was given to a word. Meltzer et al. (2013) and Kielar et al. (2016) employed tasks requiring sentence comprehension and a button-press response. In all three studies, alpha-beta (Meltzer et al., Kielar et al.) or beta (Spironelli et al.) power modulations, indexing language function, differed between patients and controls with respect to the spatial distribution of the oscillations. To the best of our knowledge, our study is the first to employ a task combining sentence comprehension and word production and to combine electrophysiological measures and an index of interhemispheric structural connectivity. Corroborating previous literature, our findings suggest that incorporating direct measures of neural activity into investigations of neuroplasticity provides an important neural marker to help predict recovery (see for discussion Reid et al. 2016), assess the progress/success of neurorehabilitation and delineate oscillatory targets for neuromodulation (Thiel et al. 2013).

In conclusion, we show a causal link between context-driven word retrieval and alpha-beta desynchronization in the left temporal-inferior parietal lobe. Moreover, we provide novel evidence that the right hemisphere can perform similar neuronal computations as the left hemisphere, as evidenced by a similar spectro-temporal neurophysiological signature of context-driven word retrieval in both hemispheres. Finally, interhemispheric posterior white-matter connections may be important to enable the right hemisphere to compensate for the loss of function following left hemisphere brain injury.

## Acknowledgements

The authors are grateful for the patients and their families, as well as for the other volunteer participants for taking part in this study. We would like to thank Donatella Scabini and Brian Curran for patient delineation, Brian Curran, Clay Clayworth and Callum Dewar for lesion reconstruction, Amber Moncrief and Selvi Paulraj for helping design the materials, Paige Mumford and Laura Agee for help with audio recordings, Colin Hoy for help with image processing, and the members of the Center for Aphasia and Related Disorders at the VAHCS in Martinez, CA, for neuropsychological testing and invaluable feedback on the text. The authors declare no conflict of interest.

